# Alterations in Connectome Dynamics in Autism Spectrum Disorder: A Harmonized Mega- and Meta-Analysis Study Using the ABIDE Dataset

**DOI:** 10.1101/2021.10.03.462909

**Authors:** Yapei Xie, Zhilei Xu, Mingrui Xia, Jin Liu, Xiaojing Shou, Zaixu Cui, Xuhong Liao, Yong He

## Abstract

**BACKGROUND:** Neuroimaging studies have reported functional connectome aberrancies in autism spectrum disorder (ASD). However, the time-varying patterns of connectome topology in ASD individuals and the connection between these patterns and gene expression profiles remain unknown.

**METHODS:** To investigate case-control differences in dynamic connectome topology, we conducted mega- and meta-analyses of resting-state functional magnetic resonance imaging data of 939 participants (440 ASD patients and 499 healthy controls, all males) from 18 independent sites, selected from the ABIDE (Autism Brain Imaging Data Exchange) dataset. Functional data was preprocessed and analyzed using harmonized protocols, and brain module dynamics was assessed using a multilayer network model. We further leveraged postmortem brain-wide gene expression data to identify transcriptomic signatures associated with ASD-related alterations in brain dynamics.

**RESULTS:** Compared to healthy controls, ASD individuals exhibited a higher global mean and lower standard deviation of whole-brain module dynamics, indicating an unstable and less regionally differentiated pattern. More specifically, ASD individuals showed higher module switching, primarily in the medial prefrontal cortex, posterior cingulate gyrus, and angular gyrus, and lower switching in the visual regions. These alterations in brain dynamics were predictive of social impairments in ASD individuals and were linked with expression profiles of genes primarily involved in the regulation of neurotransmitter transport and secretion, as well as with previously identified autism-related genes.

**CONCLUSIONS:** This study is the first to identify consistent alterations in brain network dynamics in ASD and the transcriptomic signatures related to those alterations, furthering insights into the biological basis behind this disorder.

## INTRODUCTION

Autism spectrum disorder (ASD) is a highly heritable neurodevelopmental disorder characterized by persistent impairments in social communication and the presence of restricted and repetitive patterns of behavior (1, 2). Contemporary views of ASD conceptualize it as a connectome dysfunction syndrome (3, 4), manifesting as aberrant functional connectivity in the brain, especially in the default-mode network (DMN) (5-8). ASD-related aberrancies in the brain connectome are linked with individual clinical symptoms (8-10) and impairments in cognitive ability (11, 12). These studies have provided insights into understanding the biological underpinnings of ASD from a network perspective.

However, despite its importance for understanding the disorder, previous functional connectome studies on ASD have focused primarily on the static (i.e., time-invariant) connectivity patterns, largely ignoring the temporal characteristic of brain networks. The human brain can be thought of as a highly dynamic networked system that exhibits connectivity reconfigurations over time (13, 14). These dynamic reconfigurations are essential for efficient inter-module communication (15), flexible cognitive functions (16, 17), and rapid response to external environment (18). Although several prior studies have reported alterations in brain connectome dynamics in ASD, such as increased connectivity variability (19-22) and fewer transitions between connectivity states (23-25), the topological features of dynamic brain networks in ASD remain understudied. Investigating the temporally fluctuating patterns in ASD brain network topology, in particular the properties of modular switching, will advance our understanding of how dynamic interactions of different network components underpin cognitive dysfunction and clinical symptoms in patients (26). Thus, the time-varying pattern of functional connectome topology in ASD is a pertinent area that warrants further research.

Genetic factors are considered to be a predominant cause of ASD (1). Former twin and family studies have confirmed the prominent heritability of ASD and ASD-associated traits (27). An increasing number of susceptibility genes have been identified, such as common variants (28) and rare, de novo variants (29). Recently, a large-scale exome sequencing study of ASD identified over 100 putative ASD-associated risk genes, the majority of which are neuronally expressed (30). RNA microarray and sequencing studies of post-mortem ASD brain samples also demonstrated transcriptionally altered genes and affected pathways (31). A very recent study suggests that the spatial layout of network module dynamics in healthy brains is linked to the expression level of genes associated with potassium ion channel activity regulation and mitochondria (32). Thus, we speculate that the alteration of brain network dynamics in ASD is related to the expression profile of previously identified autism-related genes.

To address these questions, we conducted the first mega- and meta-analyses for the identification of significant alterations in connectome dynamics in ASD. We used resting-state functional MRI data (rsfMRI) from 939 participants selected from 18 independent sites (33, 34) and employed a multilayer network model (35) to characterize the topological dynamics of the functional connectome. The mega- and meta-analyses were performed separately, using harmonized image processing and network analysis protocols. Finally, we conducted a partial least squares regression analysis to determine the link between abnormal network dynamics and transcriptional profiles. We hypothesized that: i) ASD patients would show significant alterations in brain connectome dynamics as compared to healthy controls (HCs), in particular in the DMN regions; and ii) these alterations in brain dynamics would be associated with individual social impairments in patients and the expression profiles of genes that were enriched for previously published ASD-related gene sets.

## METHODS AND MATERIALS

### Data Sets

We selected rsfMRI data from 440 ASD individuals and 499 HCs (all males, age range: 5-35 years old, collected at 18 independent sites) (Figure S1) from the publicly available Autism Brain Imaging Data Exchange (ABIDE) I and ABIDE II datasets (33, 34) (http://fcon_1000.projects.nitrc.org/indi/abide/) after screening against strict criteria (see Supplement).

### Data Preprocessing

All rsfMRI data were preprocessed with a standardized and harmonized pipeline using the GRETNA package (36). This process involved removing the first 10-second volumes, slice timing correction, realignment, spatial normalization to standard space, spatial smoothing with a Gaussian kernel (full width at half maximum = 6 mm), linear detrending, nuisance regression (for the following nuisance regressors: Friston’s 24 head-motion parameters, cerebrospinal fluid, white matter, and global signals), and temporal filtering (0.01-0.1 Hz). Given that rsfMRI scanning duration has been different between sites, we took data from a fixed scanning length (i.e., the first 5-minute time course) for connectome construction.

### Constructing Dynamic Brain Connectomes

For each individual, dynamic brain connectomes were generated using a sliding window approach (14, 37). Specifically, network nodes were defined as 512 regions of interest with uniform areas obtained from a random parcellation (38). Within each time window, we estimated the inter-node functional connections by calculating the Pearson’s correlation coefficient between nodal time courses. Here, the window length was set as 60 seconds and the sliding step was set as one repetition time. Finally, we obtained weighted dynamic connectomes by applying a network threshold with a fixed density (density = 15%) to reduce the influence of weak or spurious connectivities (16).

### Tracking Dynamic Modular Structures

We utilized a multilayer network model (35) to identify the time-varying features of connectome topology (Figure 1A). This model incorporates connectivity information from adjacent windows and assumes temporal continuity of modular configurations. Specifically, we conducted a multilayer-variant Louvain algorithm (http://netwiki.amath.unc.edu/GenLouvain) to identify the optimal modular architecture by maximizing the modularity index, Q (range: 0-1), which denotes the extent of segregation between network modules. Then, we computed the modular variability (16) of each brain node to quantify how individual nodes dynamically switched their modular affiliations over time (see Supplement). The larger the modular variability, the more flexibly a brain node switches between modules.

**Figure 1.**
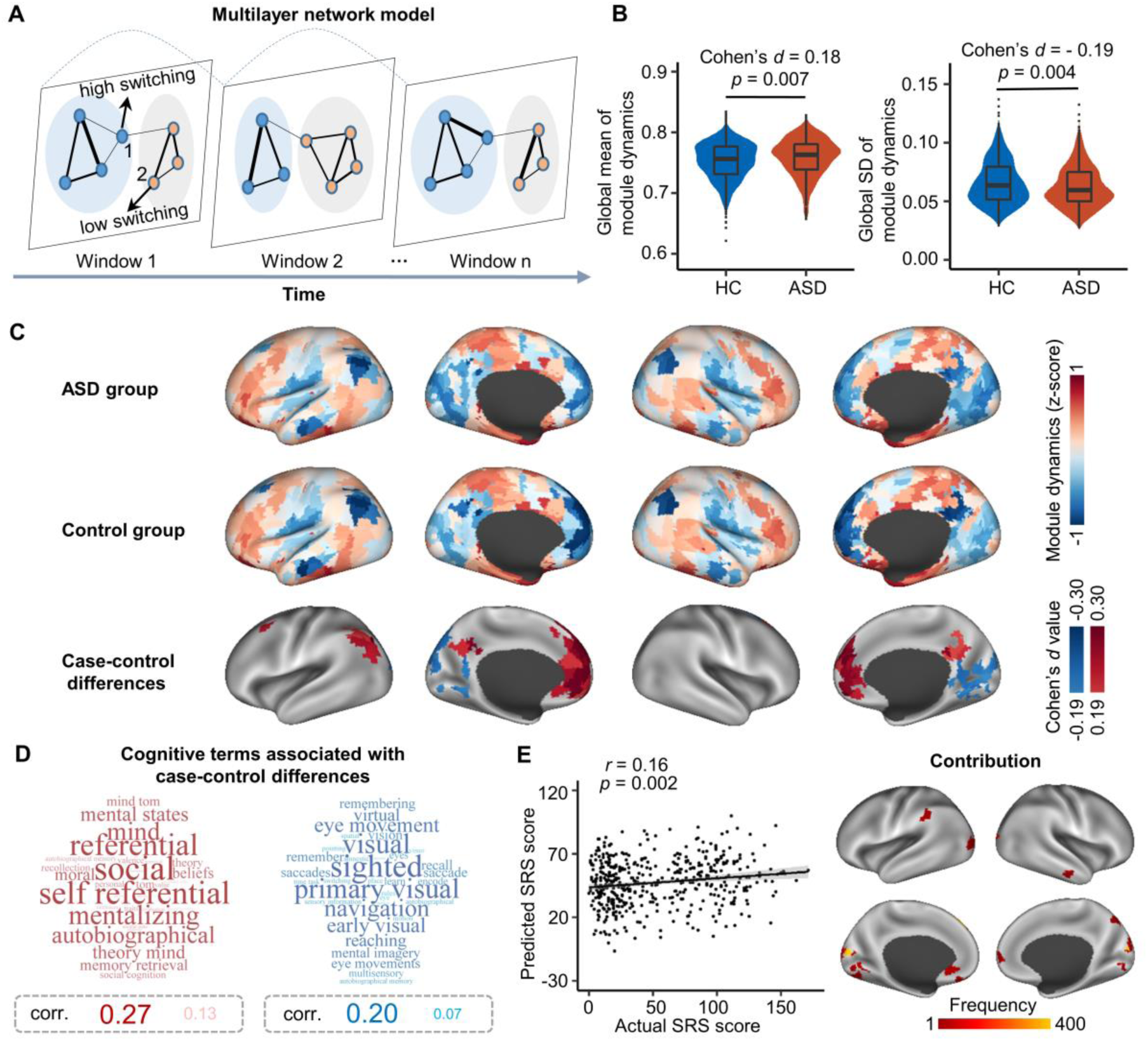
Mega-analysis of case-control differences in module dynamics. (A) Schematic overview of brain module switching within a multilayer network model. Each node not only connects to nodes in the same window but also connects to itself in the two temporally adjacent windows. Node colors denote nodal module affiliations. Modular switching is determined by modular variability, which reflects the module dynamics. Node 1 shows a high modular variability while node 2 shows a low modular variability. (B) Mega-analysis of case-control differences in the mean value and the standard deviation of modular variability at the global level. (C) Mega-analysis of case-control differences in module dynamics at the nodal level. The upper and middle panels show the group-level modular variability maps for each population. The lower panel shows regions with significant case-control differences in modular variability, corrected for multiple comparisons (FDR-corrected *p* < .05, corresponding to *p*_*uncorr*_ < .004). (D) Cognitive terms associated with the regions showing significant case-control differences. The red and blue word clouds respectively represent cognitive terms associated with regions showing significantly higher and lower module dynamics in the ASD group. Font size has been scaled to reflect the correlation value for each cognitive term. (E) Prediction of individual SRS scores based on modular variability maps using support vector regression. The scatterplot displays the correlation between actual and predicted SRS scores. Each dot corresponds to one instance of leave-one-out cross-validation. The brain map displays regional contribution to the prediction, which was defined as the frequency that each region was selected as a feature in the leave-one-out cross-validation. SD, standard deviation; ASD, autism spectrum disorder; SRS, Social Responsive Scale.

### Case-Control Comparison Analysis

#### Mega-Analysis

To examine case-control differences in module dynamics, we performed a mega-analysis by pooling individual modular variability maps across all sites. Prior to performing the analysis, we applied a ComBat harmonization (39-41) to the modular variability maps to correct for site effects. We then estimated group differences in modular variability at both global (whole-brain mean and standard deviation) and nodal levels using a semi-parametric Generalized Additive Model (GAM) (42) with restricted maximum likelihood as the smoothing parameter:

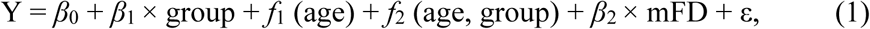

where Y denotes the measure of modular variability. The age and age-by-group interaction effects were controlled for by introducing two smooth functions (i.e., *f*_1_, *f*_2_) as non-parametric terms. This allows for flexible assessment of the nonlinear relationship without pre-emptively assigning a prior shape. Mean framewise displacement (43) was also included as a covariate to control for head motion. Multiple comparisons were corrected for by applying the false discovery rate (FDR) method (44). Cohen’s *d* values, representing the effect size of the group comparisons, were computed from the *t* statistic of the group term. The GAM was computed using the mgcv package (https://cran.rproject.org/web/packages/mgcv/index.html). To further decode the cognitive implications of the brain nodes exhibiting ASD-related alterations in connectome dynamics, we performed a functional meta-analysis using the Neurosynth database (45) (see Supplement).

#### Meta-Analysis

To assess the robustness of the observed case-control differences in the mega-analysis, we also undertook a harmonized meta-analysis. Briefly, for each site we conducted the GAM (Eq. (1)) to examine site-specific group differences in modular dynamics at both global and nodal levels. Then, we obtained the meta-analytic Cohen’s *d* values of these measures using an inverse variance-weighted random effect meta-analysis model in the metafor package (version 3.0.2; https://cran.r-project.org/web/packages/metafor/index.html).

### Prediction of Social Impairments Using Connectome Dynamics

We further evaluated whether brain module dynamics were predictive of individual social impairments observed in ASD. To quantify the degree of social impairments, we referred to scoring against the Social Responsiveness Scale (SRS), which provides a dimensional characterization of the severity of social impairments related to ASD. We trained a support vector regression model to estimate each participant’s SRS score based on the whole-brain modular variability maps. The leave-one-out cross validation (LOOCV) was used to estimate the accuracy of our predictions. In each LOOCV fold, we included the feature selection, model learning, and testing. Nodal contribution to the prediction was defined as the frequency that each node was selected as a feature during the LOOCV (see Supplement). This analysis was performed using the LIBSVM toolbox (46).

### Association between Alterations in Connectome Dynamics and Gene Expression Profiles

#### Estimation of Gene Expression in Brain Nodes

We utilized the genome expression data from five male post-mortem human brains from the Allen Human Brain Atlas dataset (47) to identify genes associated with ASD-related alternations in connectome dynamics. Gene expression levels from the left hemisphere was used here, since right hemisphere data was available from only two donors. The microarray data was preprocessed using a state-of-the-art analysis pipeline (48) and spatially matched with 222 brain nodes (see Supplement). This resulted in a 222 × 10,145 matrix, denoting the expression of 10,145 genes across 222 nodes.

#### Spatial Correlation with Gene Expression Profiles

Given the high similarity of results obtained from both the mega- and meta-analyses in respect of ASD-related alterations in brain module dynamics (see Results), we performed the connectome-transcriptome association analysis based on the group difference map (i.e., Cohen’s *d* values) from the mega-analysis. Specifically, we used a partial least squares (PLS) regression to identify the weighted linear combinations (i.e., components) of expression patterns for all 10,145 genes, which were correlated with ASD-related alterations in connectome dynamics. The statistical significance of the variance explained by the PLS components were tested using a permutation analysis (n = 10,000) in which spatial autocorrelation was corrected for (49). For each PLS component map, we calculated the spatial similarity between the weighted gene expressions and the Cohen’s *d* values in the group difference map using the Pearson’s correlation. The significance of the correlation was tested again using a permutation analysis (n = 10,000) in which spatial autocorrelation was corrected for (49). Finally, the PLS weight of each gene was transformed into a Z-score value by dividing the weight by the standard deviation of the corresponding weights derived from 1,000 instances of bootstrapping (resampling with replacement of 222 nodes). We then ranked all genes according to their Z-score weights to the PLS components.

#### Enrichment Analysis

To explore the functional significance of the associated genes, we first conducted separate searches for Gene Ontology terms that were enriched at the top (strong positive correlation) and bottom (strong negative correlation) of the ranked gene list, by employing the widely used online tool GOrilla (50) (http://cbl-gorilla.cs.technion.ac.il/) (see Supplement). All three ontology classes, including biological process, cellular component, and molecular function, were considered.

Next, we performed a gene set enrichment analysis (51) on the whole gene list (i.e., the ordered set of 10,145 genes) to assess whether ASD-related gene sets identified in previous studies were overrepresented in the most strongly correlated genes identified in our ordered list. Specifically, we considered six different classes of ASD-related gene sets (Table S1), including i) gene set 1, ASD-related genes from a summary of multiple datasets (52); ii) gene set 2, ASD risk genes from a large-scale exome sequencing study (30); iii) gene set 3, ASD-associated common genetic variants from a genome-wide association meta-analysis study (28); iv) gene set 4, ASD associated rare, de novo variants from a study integrating copy number variants and sequencing data (29); v) gene set 5, genes upregulated in the ASD cortex from a post-mortem genome-wide transcriptome study (31); and vi) gene set 6, genes downregulated in the ASD cortex from a post-mortem genome-wide transcriptome study (31). For the purposes of comparison, we also included one gene set that was associated with non-mental health diseases (i.e., gene set 7) (52). The enrichment analysis was performed using the clusterProfiler package, version 3.14.3 (53) (https://bioconductor.org/packages/release/bioc/html/clusterProfiler.html).

### Power Estimation

We estimated the minimal effect size (i.e., Cohen’s d) observable for case-control differences between 440 ASD individuals and 499 HCs using G*Power, version 3.1.9.4. At a significant threshold of .05 (two-tailed) and a minimum desired power level of .8, we have the statistical power to observe Cohen’s d greater than .18.

### Validation Analysis

We validated our main findings by considering five potential confounding factors, including head motion, window length, intelligence quotient, imaging sites, and age range (see Supplement).

## RESULTS

### Demographic Characteristics

Table 1 summarizes the demographic and clinical information of the participants. No significant difference in age was found between the ASD and HC groups (*p* = .84). The ASD group showed lower IQ scores (*p* = 4.1 × 10^−9^) and higher SRS scores than the HC group (*p* = 4.5 × 10^−133^).

**Table 1.** Demographics and clinical characteristics of participants

| | ASD ($n = 440$) | HC ($n = 499$) | $p$ value |
| --- | --- | --- | --- |
| Age (Years) | 5.3-34.5 ($14.3 \pm 5.4$) | 5.9-34.1 ($14.4 \pm 5.4$) | .84 |
| FIQ | 71-149 ($107.1 \pm 16.6$) | 73-148 ($112.8 \pm 12.9$) | $4.1 \times 10^{-9}$ |
| ADOS-2 Total ($n = 208$) | 2-26 ($11.7 \pm 4.2$) | NA | |
| ADOS-2 Severity ($n = 208$) | 1-10 ($6.7 \pm 2.0$) | NA | |
| ADOS-2 Social ($n = 206$) | 1-20 ($8.7 \pm 3.6$) | NA | |
| ADOS-2 RRB ($n = 209$) | 0-7 ($3.0 \pm 1.6$) | NA | |
| ADI-R Social ($n = 322$) | 4-30 ($19.3 \pm 5.4$) | NA | |
| ADI-R Verbal ($n = 322$) | 4-25 ($15.6 \pm 4.4$) | NA | |
| ADI-R RRB ($n = 322$) | 0-12 ($5.7 \pm 2.6$) | NA | |
| SRS (ASD/HC: 220/253) | 16-171 ($90.5 \pm 28.7$) | 0-85 ($19.8 \pm 13.5$) | $4.5 \times 10^{-133}$ |

Note: Male participants were selected from 18 of the sites that contributed to the ABIDE I and ABIDE II datasets using stringent quality control criteria. For each clinical measure, we only considered the module/version with the largest sample of participants available. The $p$ value was obtained using a two-sample two-tailed $t$ -test. ASD, autism spectrum disorder; HC, healthy control; FIQ, full-scale IQ; ADOS-2, Autism Diagnostic Observation Schedule, Second Edition; RRB, restricted and repetitive behavior; ADI-R, Autism Diagnostic Interview–Revised; SRS, Social Responsiveness Scale; NA, not applicable.

### Alterations of Module Dynamics in ASD Connectomes

#### Mega-Analysis

At the global level, no significant group difference was found in brain network modularity (*p* = .32). However, the ASD group showed a higher mean value (Cohen’s *d* = .18, *p* = .007) and a lower standard deviation (Cohen’s *d* = -.19, *p* = .004) in whole-brain modular variability than the HC group (Figure 1B). This suggests that global brain dynamics in ASD tends to be more unstable and regionally undifferentiated as compared to that in HCs.

At the nodal level, for both the ASD and HC groups, we observed higher modular variability primarily in the bilateral prefrontal regions and the medial temporal lobe, and lower variability mainly in the medial prefrontal and parietal regions, angular gyrus, and visual cortex (Figure 1C, upper and middle panels). This pattern is highly comparable to that shown in previous studies in healthy brains (16, 32). In comparison to the HC group, the ASD group showed higher modular variability mainly in several default-mode regions, including the medial prefrontal cortex, posterior cingulate gyrus, and angular gyrus (Cohen’s *d* ranging from .19 to .33), and lower variability primarily in the visual cortex (Cohen’s *d* ranging from -.24 to -.19) (FDR corrected *p* < .05) (Figure 1C, lower panel; age and age-by-group interaction effects are described in Figure S2 and Figure S3 of the Supplement).

Using the NeuroSynth meta-analytic database (45), we found that the regions showing higher modular dynamics in ASD were mainly associated with social function and internally oriented processes, while those showing lower modular dynamics were involved in visual-related tasks (Figure 1D).

#### Meta-Analysis

At the global level, our harmonized meta-analysis revealed that, compared to the control group, the ASD group showed a higher global mean (Cohen’s *d* = .15, *p* = .01) and a lower standard deviation (Cohen’s *d* = -.23, *p* = .001) in whole-brain modular variability (Figure 2A and 2B). At the nodal level, the case-control difference pattern was remarkably similar to that derived from the mega-analysis (spatial similarity: *r* = .96, *p* < .0001 after correcting for spatial autocorrelation) (Figure 2C and Figure S4). The meta-analysis also revealed significant group differences in default-mode (Cohen’s *d* ranging from .19 to .34) and visual regions (Cohen’s *d* ranging from -.34 to -.17) (FDR corrected *p* < .05) (Figure 2D).

**Figure 2.**
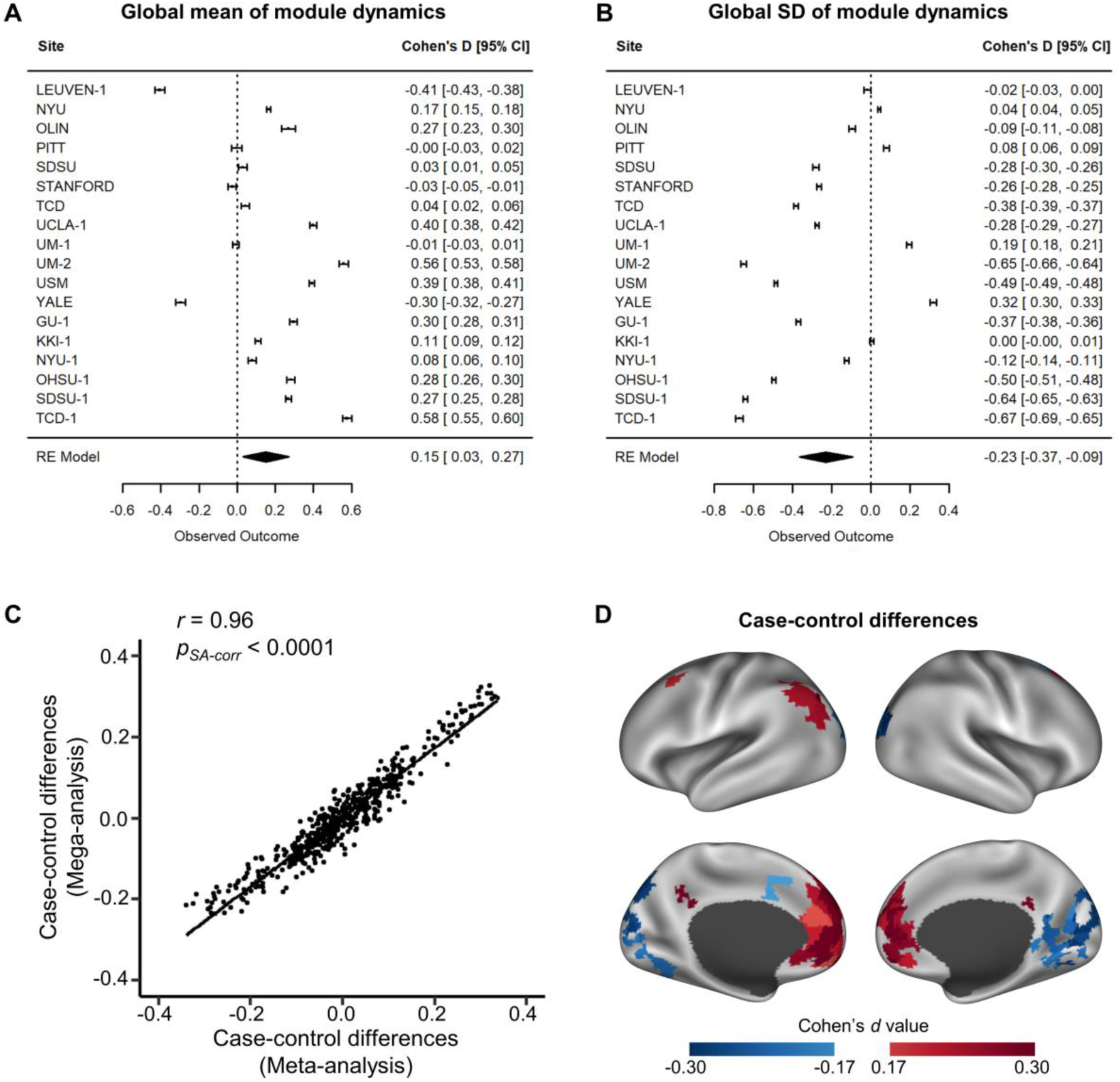
Meta-analysis of case-control differences in module dynamics. (A) Forest plot of Cohen’s *d* effect sizes for case-control differences in the global mean of modular variability. Each row shows the Cohen’s *d* effect size and the confidence intervals for each site. The meta-analysis results are displayed at the bottom with the combined effect and the confidence interval plotted as a diamond. (B) Forest plot of Cohen’s *d* effect sizes for case-control differences in the standard deviation of modular variability. (C) Spatial similarity between case-control difference maps obtained from the mega- and meta-analyses. Each dot represents a brain node. The significance level of the spatial association was corrected for spatial autocorrelation (49). (D) Meta-analysis of case-control differences at the nodal level. Significance levels of case-control differences in modular variability have been corrected for multiple comparisons (FDR-corrected *p* < .05, corresponding to *p*_*uncorr*_ < .0056). SD, standard deviation; FDR, false discovery rate.

### Predicting the Severity of Social Impairments Based on Brain Module Dynamics

Using individual modular dynamics patterns as the feature in the support vector regression model, we found that brain dynamics was a significant predictor of SRS scores (*r* = .16, *p*_*perm*_ = .002) (Figure 1E). Brain nodes making the largest contribution to SRS score prediction were mainly located in the medial prefrontal cortex and the visual cortex (Figure 1E). These regions largely overlapped with those showing case-control differences in brain modular dynamics.

### Association between Alterations in Connectome Dynamics and Gene Expression Profiles

#### Partial Least Squares Regression Analysis

We assessed the spatial association between ASD-related dynamics alterations and nodal gene expression profiles (Figure 3A). The weighted gene expression pattern of the first PLS component (PLS1) accounted for the greatest spatial variance (20.5%, Figure 3B) in modular dynamics in the case-control difference map (*p* = .05, corrected for spatial autocorrelation). The PLS1 score map was spatially correlated with the group difference map (*r* = .45, *p* = .0043, corrected for spatial autocorrelation) (Figure 3C and 3D).

**Figure 3.**
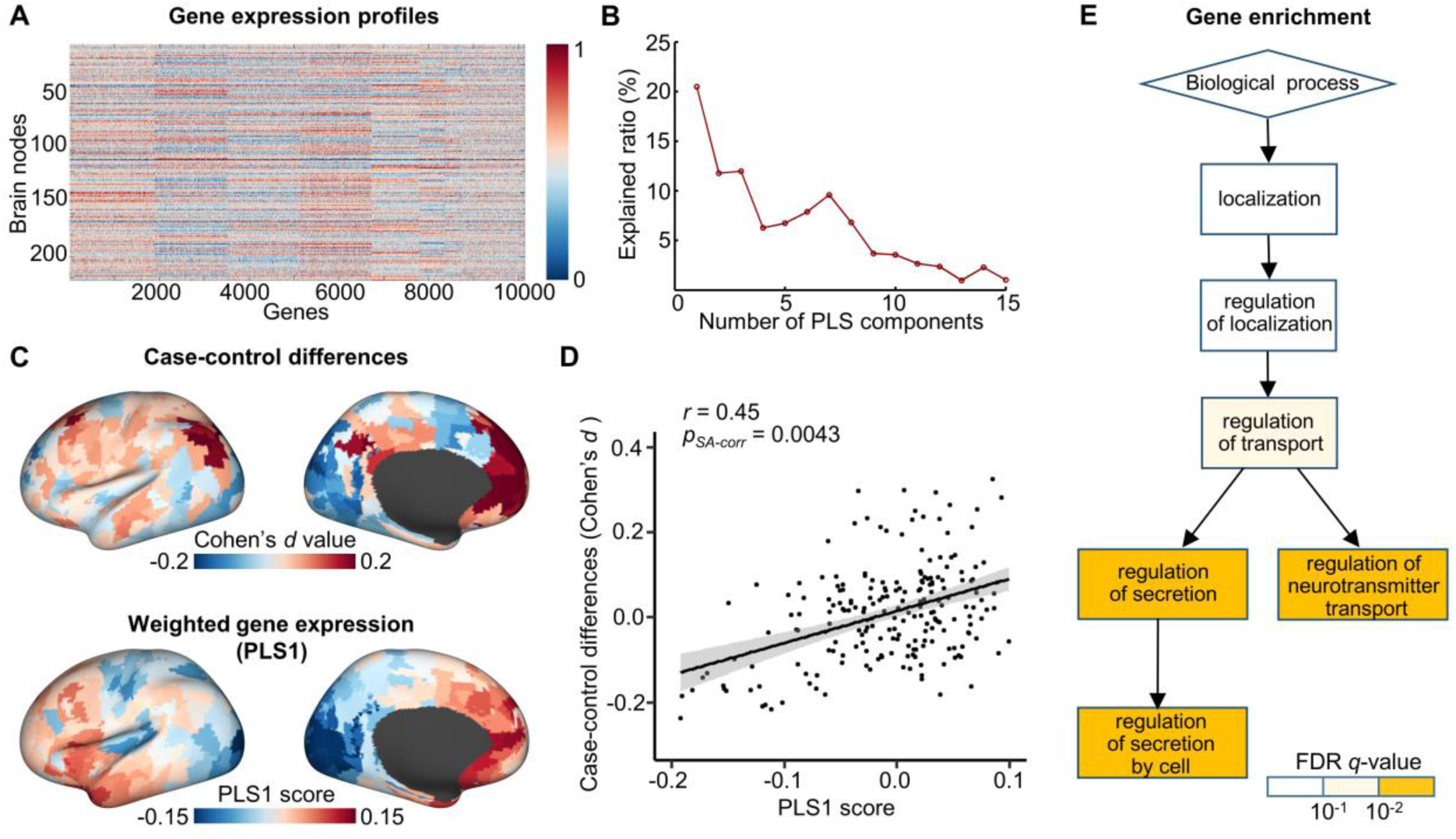
Association between ASD-related alterations in module dynamics and gene expression profiles. (A) Gene expression profiles across brain nodes. Each row denotes the gene expression for each gene at a given brain node. (B) Explained ratios for the first 15 components obtained from the partial least squares (PLS) regression analysis. Each component denotes a weighted linear combination of the expressions of all genes. (C) Spatial patterns showing the mega-analysis case-control differences in modular variability and the PLS1 scores in the left hemisphere (unthresholded). (D) Spatial association between case-control differences in modular variability and PLS1 scores. Each dot represents a brain node. The significance level of the spatial association has been corrected for spatial autocorrelation (49). (E) Significant enrichment of Gene Ontology terms associated with biological processes was observed for PLS1. Color denotes the *q*-values for the significantly enriched Gene Ontology terms. PLS, partial least squares.

#### Enrichment Analysis

We identified three biological process terms significantly enriched at the top of the gene list, including the regulation of secretion by cell, the regulation of neurotransmitter transport, and the regulation of secretion (FDR corrected *p* < .01) (Figure 3E). Interestingly, all three of these terms were related to the regulation of transport. We did not find any significant enrichment of Gene Ontology terms at the bottom of the gene list. Moreover, no significant enrichment of molecular function and cellular components was observed.

We further conducted a gene set enrichment analysis to examine whether six classes of previously reported ASD-related genes were significantly enriched at the top or bottom of our ordered gene list (Figure 4A). We found that ASD-related genes from a summary of multiple databases (i.e., gene set 1) were significantly enriched at the top of our gene list (normalized enrichment score (*NES*) = 1.27, *p*_*adjust*_ = .035, FDR corrected, hereafter the same) (Figure 4B). The ASD risk genes identified from a large-scale exome sequencing study (i.e., gene set 2) exhibited a significant enrichment at the bottom of our gene list (*NES* = -1.48, *p*_*adjust*_ = .035) (Figure 4C). We also observed that ASD-related common genetic variants (i.e., gene set 3) were significantly enriched at the bottom of our gene list (*NES* = -1.70, *p*_*adjust*_ = .028) (Figure 4D), while ASD-related rare, de novo variants (i.e., gene set 4) exhibited only marginally significant enrichment (*NES* = -1.39, *p*_*adjust*_ = .069) at the bottom of our gene list (Figure 4E). Moreover, genes upregulated and downregulated in the post-mortem ASD cortex (i.e., gene sets 5 and 6) were significantly enriched at the top and bottom of our gene list, respectively (upregulated: *NES* = 1.41, *p*_*adjust*_ = .009; downregulated: *NES* = -1.67, *p*_*adjust*_ = .002) (Figure 4F and Figure 4G). As a control dataset, the gene set comprising genes associated with non-mental-health diseases (i.e., gene set 7) was not significantly enriched at the gene list (*NES* = -.99, *p*_*adjust*_ = .51) (Figure 4H).

**Figure 4.**
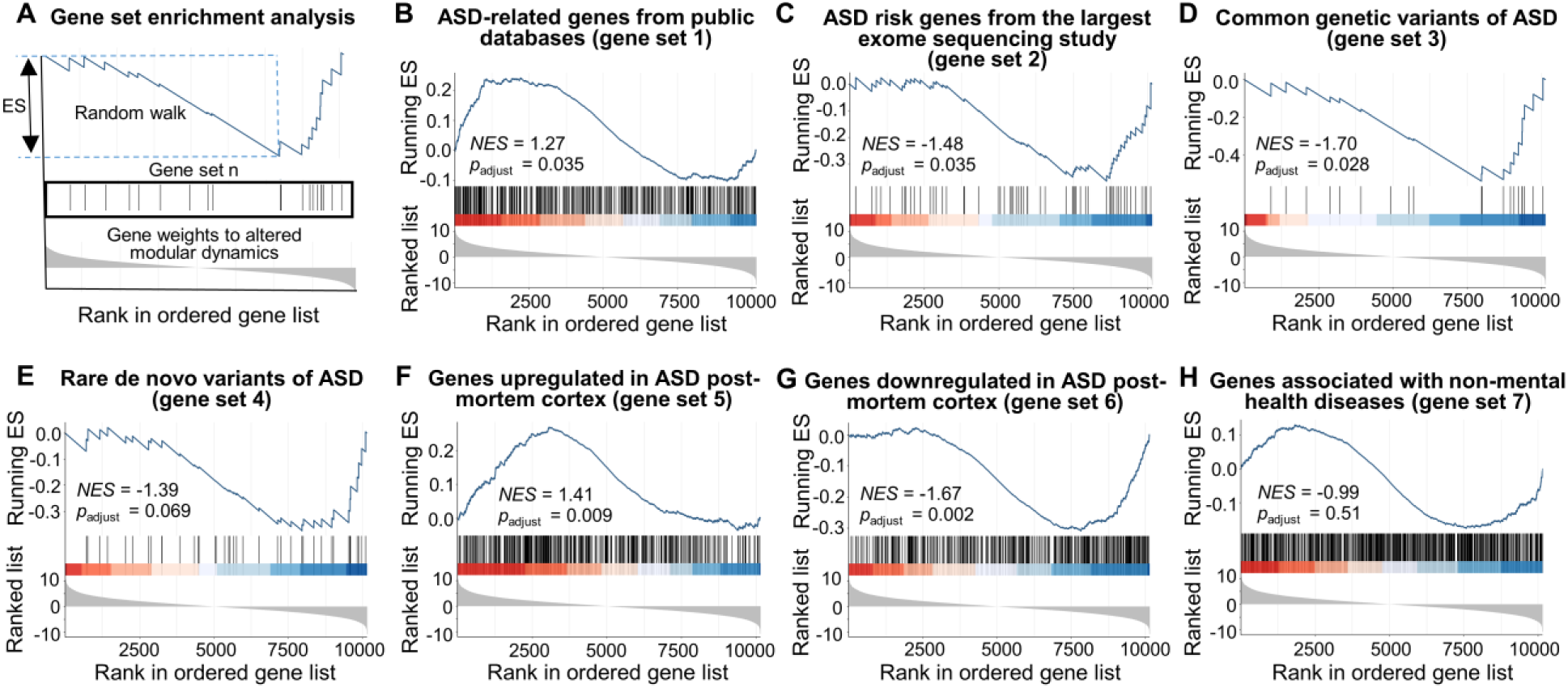
Gene set enrichment analysis of genes associated with ASD-related alterations in module dynamics. (A) An overview of the gene set enrichment analysis. The solid line denotes the running enrichment score (*ES*) along the ordered gene list, which increases when a gene is included in the gene set of interest and decreases when a gene is not included. The vertical lines in the middle display the locations at which the members of the gene set appear in the ordered gene list. The shaded curve at the bottom denotes the value of the ranking metric (i.e., Z value for each gene’s PLS1 weights) for the genes in the ordered gene list. The *ES* captures the degree to which the gene set is overrepresented at the top or bottom of the ordered gene list, which is defined as the maximum deviation from zero of the running *ES*. Significance of the *ES* was estimated by a gene set-based permutation (10,000 times). (B) Significant enrichment of ASD-related genes from a summary of multiple databases (52). (C) Significant enrichment of ASD risk genes (30). (D) Significant enrichment of common genetic variants of ASD derived from a GWAS study (28). (E) Marginally significant enrichment of ASD-related rare, de novo variants. (F) Significant enrichment of genes upregulated in the post-mortem ASD cortex (31). (G) Significant enrichment of genes downregulated in the post-mortem ASD cortex (31). (H) Non-significant enrichment of the gene set associated with non-mental health diseases (52). After the gene set enrichment analysis was performed for all gene sets, the *ES* for each gene set was normalized to the *NES* to account for the gene set size, and multiple comparisons with the seven gene sets were corrected for using the false discovery rate method. ASD, autism spectrum disorder; PLS, partial least squares; *ES*, enrichment score; *NES*: normalized enrichment score; GWAS, genome-wide association study.

### Validation Results

When assessing the potential influence of five confounding factors, we found that ASD-related, significant alterations in brain modular dynamics remained highly similar to our main results (Dice index ranging from .77 to .95) (Table S2 and Figure S5, see Supplement). This suggests that our results were robust and not affected by methodological variations.

## DISCUSSION

Using a harmonized mega- and meta-analysis, this study provides the first robust demonstration of ASD-related alterations in brain modular dynamics. Our study reveals that these alterations occur primarily in the default-mode and visual regions and are associated with social impairments in patients and with the expression profiles of genes enriched for the regulation of neurotransmitter transport and secretion, as well as with previously reported autism-related genes. Together, these findings provide evidence for altered macroscopic connectome dynamics and illustrate its linkage with microscopic transcriptional profiles, advancing our knowledge of the biological mechanisms behind ASD.

### Aberrant Configuration of Dynamic Modular Architecture in ASD

Previous studies have reported on the existence of temporal modular switching in healthy brain networks (16, 37, 54, 55), which is critical for efficient integration between different functional systems (56) and for cognitive flexibility (16). Here, we demonstrate that patients with ASD exhibited abnormal, higher levels of modular switching in several default-mode regions (in particular the medial prefrontal cortex) but lower modular switching in visual areas. This reflects more frequent switching and excessive functional integration between the default-mode network and other networks over time, but reduced functional integration between the visual network and other networks. Prior studies have also reported altered dynamic connectivity in the default-mode (19, 24, 57) and visual regions (23, 58, 59) in ASD. Compared with these studies, our findings extend the existing knowledge of ASD-related abnormalities in brain dynamics from the connectivity-level to the system-level, advancing our understanding of ASD psychopathology from a dynamic connectome topology perspective.

Mounting evidence has suggested an association between social functions and the default-mode and visual regions. For instance, default-mode regions (especially the medial prefrontal cortex) have been demonstrated to be involved in social inference (60). At the same time, visual perception impairments significantly contribute to early social-emotional deficits (61) and have cascading effects on learning and social development in ASD (62). These findings hint that the alterations in connectome dynamics in the default-mode and visual regions identified here may underlie social impairments in ASD. Indeed, we observed that brain module dynamics patterns significantly predicted individual social impairments, suggesting that temporal characteristics in connectome topology may be a promising neuroimaging biomarker for ASD symptoms.

### Transcriptional Profiling of Aberrant Brain Module Dynamics

Leveraging postmortem brain-wide gene expression data from the AHBA (47), we found that ASD-related alterations in brain dynamics were closely associated with the transcriptional profiles of genes involved in the regulation of neurotransmitter transport and secretion. Extensive studies have indicated that aberrant neurotransmitter transport is a significant feature of ASD, especially aberrancies in the transport of the excitatory neurotransmitter glutamate and the inhibitory neurotransmitter gamma-aminobutyric acid (63). This suggests that brain regions with abnormal dynamics in ASD, e.g. the medial prefrontal and visual cortices, may have failed to maintain balanced excitatory-inhibitory neurotransmitter transport. Such speculations are supported by prior studies that have reported imbalanced excitatory-inhibitory synaptic transmission in these regions (64-66). Accordingly, our findings may provide support for the existing excitation/inhibition imbalance theory in ASD (66-68) by revealing a transport regulation-specific connectome-transcriptome association.

We also demonstrated that previously reported autism-related genes were significantly enriched among the genes that we found to be most strongly correlated (both positively and negatively) with alterations in modular dynamics, indicating that different classes of genes contribute to the alterations in module dynamics in ASD. Specifically, compared with rare, de novo variants (gene set 4) (29), ASD-related common genetic variants showed a more substantial influence on module dynamics, which was manifested as a significant enrichment of the gene set. This finding is consistent with a recent study regarding cortical volume and transcriptome association in ASD (69), and could be due to common variants explaining a larger proportion of heritable variance in ASD compared to rare, de novo variants (70). Moreover, we found that genes upregulated and downregulated in the post-mortem ASD cortex were respectively overrepresented at either end of our gene list. As the top and bottom weighted genes in the ranked list showed different correlations (i.e., positive and negative) with ASD-related alterations in module dynamics, we speculate that these two categories of dysregulated genes in ASD are likely to affect brain network dynamics in different ways. Several previous studies have also shown the association between abnormal brain morphology (e.g., cortical thickness and volume) and the downregulated genes in ASD, but the same association was not found for the upregulated genes (69, 71). Combining these findings, the implication is that different imaging phenotypes in ASD may show common and specific genetic factors.

Notably, we did not find any significant overrepresentation of genes in the gene set associated with non-mental health diseases (gene set 7) (52), suggesting that the genes identified as being strongly correlated were mental disorder-specific. Together, the results from our connectome-transcriptome association analysis provides a meaningful link between the ASD-related alterations in macroscopic brain dynamics and microscopic molecular signatures.

### Limitations and Future Work

Several issues should be taken into consideration when interpreting these results. First, only male participants were included in this study, given the high prevalence of ASD in males and the small sample of female participants in the ABIDE database. Whether the findings can be generalized to the female population still needs further investigation. Second, gene expression data used here was derived from five healthy brains. It will therefore be promising to explore the connectome-transcriptome association using ASD gene expression data when the relevant data becomes available. Third, we did not exclude medicated ASD individuals, given the limited medication information available from the ABIDE database. Whether and how medication affects the alterations in module dynamics remains for further exploration. Finally, as a psychiatric disorder, ASD often features high comorbidity and may exhibit alterations in brain function and genetic factors that are also present in other brain disorders (72, 73). Revealing ASD-specific genes and dynamic connectome alterations will be important for understanding the biological basis of this disorder.

## ACKNOWLEDGMENTS

This study was supported by the National Natural Science Foundation of China (Nos. 82021004, 81971690, 31830034, 81620108016, 81801779, and 11835003), the Changjiang Scholar Professorship Award (T2015027), the Beijing Brain Initiative of the Beijing Municipal Science & Technology Commission (Z181100001518003), and the Fundamental Research Funds for Central Universities (2019NTST24).

## DISCLOSURES

The authors report no biomedical financial interests or potential conflicts of interest.

## Supplementary Information

### Supplementary Methods

#### Data Sets

Resting-state functional MRI (rsfMRI) data covering 2,156 participants was initially obtained from the ABIDE I (17 sites, ASD/HC: 539/573, 7-64 years) and ABIDE II (19 sites, ASD/HC: 487/557, 5-64 years) databases (http://fcon_1000.projects.nitrc.org/indi/abide/). We screened this initial dataset using strict inclusive criteria and only included scans that: i) were taken from male participants, as 94% of participants were male; ii) were from participants who did not show any severe structure damage in the T1 images; iii) consisted of single-band fMRI data, as only two sites used multiband scanning profiles; iv) had a scan length of at least 5 minutes; v) had maximal head motion of less than 5 mm and 5 degrees, and mean framewise displacement (FD) (1) of less than 0.5 mm; vi) had nearly full-brain coverage and successful spatial normalization; vii) were taken from participants with a full scale IQ higher than 70. Finally, we included only scans from participants under 35 years old, due to the limited number of participants older than 35. We also excluded data from sites that had fewer than 10 participants remained in either ASD or control group after the above screening had been applied. Thus, after all selection criteria had been applied, we obtained a dataset consisting of scans from 939 participants (ASD/HC, 440/499) taken from 18 different sites (Figure S1).

#### Tracking Dynamic Modular Structures

We utilized a multilayer network model (2) to identify the window dependent (i.e., time-varying) modular architecture. This algorithm incorporates connectivity information from temporally adjacent two windows and assumes temporal continuity of the modular architecture across windows. Specifically, the window-dependent functional network was treated as a multilayer network, in which each node not only connected to nodes within the same layer (i.e., window) but also connected to the identical nodes in two temporally adjacent layers (i.e., one back and one forward). The optimal modular architecture was identified by maximizing the multilayer modular index, Q, which is defined as (2)

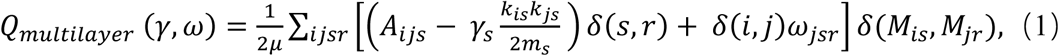

where *i* and *j* are node labels and *s* and *r* are layer labels.

Specifically, *μ*denotes the total degree of the multilayer network, *A*_*ijs*_ denotes the connectivity strength between nodes *i* and *j* in layer *s, k*_*is*_ denotes the degree of node *i* in layer *s, m*_*s*_ denotes the total degree of layer *s, k*_*is*_*k*_*js*_/2*m*_*s*_ denotes probability expected by chance of a connection between node *i* and node *j* in layer *s*. The function *δ*(*x, y*) equals 1 if *x* equals *y*, and equals 0 otherwise. The variable *γ*_*s*_ is the topological resolution parameter in layer *s*, which determines the detected module size. The larger the value of *γ*_*s*_, the smaller the detected module size. Parameter *ω*_*jsr*_ is a temporal coupling parameter, denoting the strength of inter-layer coupling for node *j* between layers *r* and *s*. Here, we used the commonly used default values of *γ* = 1 and *ω* = 1 (3, 4).

Due to the heuristic nature of modularity optimization based on the multilayer-variant Louvain algorithm (5), we repeated the multilayer modular detection process 100 times. All module-relevant measures (i.e., modularity and modular variability) were calculated as the averaged value across the 100 runs. The Genlouvain MATLAB package was used to perform multilayer community detection (http://netwiki.amath.unc.edu/GenLouvain).

Given the time-varying nature of the modular architecture, we employed a measure of modular variability (6) to quantify how each brain region switched between modules over time. For example, given a node *k*, its modular variability between two sliding windows *i* and *j* is defined as

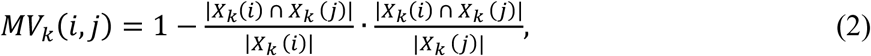

where *X*_*k*_ (*i*) and *X*_*k*_ (*j*) denote the module labels to which node *k* belongs in window *i* and *j* respectively, *X*_*k*_(*i*) *∩ X*_*k*_(*j*) denotes the set of common nodes between modules *X*_*k*_ (*i*) and *X*_*k*_ (*j*), and |*X*| denotes the node number in the node set *X*. For node *k*, its total modular variability across all windows is calculated as

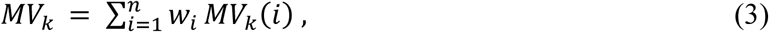

where *MV*_*k*_ (*i*) *=* Σ_*j*≠*i*_ *MV*_*k*_(*i, j*)/(*n* − 1) denotes the modular variability of node *k* between windows *i* and all other windows. In Eq. (3), we used a normalized weighed coefficient *w*_*i*_ to reduce the bias of potential outlier time windows. The coefficient *w*_*i*_ denotes the spatial similarity in the modular architecture between window *i* and all other windows, and is estimated using adjusted mutual information (7). The larger the modular variability, the more frequently a brain node switches between modules. For each node, we estimated its modular variability across all windows, and thus obtained a modular variability map for each participant.

#### Site Effect Correction in the Mega-Analysis

To reduce the influence of different sites, we applied a ComBat harmonization (8-12) to the individual modular variability maps to correct for site effects before subsequent statistical analysis. The Combat approach removes unwanted inter-site variability in imaging acquisition while preserving inter-site variability in biologically meaningful metrics. We retained age, group, mean FD, and IQ as regressors in the ComBat process to preserve biologically meaningful variability. Of note, for the nodal-level analysis, we transformed the harmonized modular variability values into z-score values to further enhance the comparability between sites. The nodal z-score values were obtained by subtracting the mean modular variability across the brain and then dividing by the corresponding standard deviation.

#### Case-Control Comparison Analysis in the Mega-Analysis

We assessed differences in module dynamics between the ASD and HC groups using a semi-parametric Generalized Additive Model (GAM) (13) with restricted maximum likelihood as the smoothing parameter. The age and age-by-group interaction effects were controlled by introducing two smooth functions for these non-parametric terms. The significance level of all three terms (i.e., group, age, and age-by-group) at the nodal level was corrected for using a false discovery rate (FDR) method (14) in which the *p* values of all three effects were pooled together (15). In the main analysis, we reported only results on the group effect of interest (i.e. case-control differences in module dynamics). The age and age-by-group effects on module dynamics are provided here as a supplement (see Figures S2 and S3).

#### Correlation between ASD-related Alterations and Cognitive Terms

To decode the cognitive implications of the divergences observed between the ASD and HC groups, we performed a functional meta-analysis using the Neurosynth database (16). We first generated a *t*-value map from the mega-analysis for ASD-related alterations, showing the brain nodes exhibiting significant differences in modular variability. Using this map, we then generated two separate t-value maps for ASD-related increases in modular variability and ASD-related decreases in modular variability, respectively. Next, for each map, we decoded the associated cognitive terms and retained the top 35 as having substantial relevance.

#### Prediction of Social Impairments Using Connectome Dynamics

We further evaluated whether the spatial pattern of nodal modular variability was predictive of individual social impairments. To quantify the degree of social impairments, we referred to scoring against the Social Responsiveness Scale (SRS), which provides a dimensional characterization of the severity of ASD-related social impairments, and which was available for the largest number of participants (i.e., 220 ASD individuals and 253 HCs).

We used a support vector regression (SVR) model to estimate each participant’s SRS score based on the whole-brain nodal modular variability values. The leave-one-out cross-validation (LOOCV) was used to estimate prediction accuracy. Specifically, in each LOOCV fold, one participant was designated as the test set, and the remaining participants were set as the training set. We first generated a general linear model for the training set to reduce the effects of age and mean FD, and applied the estimated parameters to the test set. The resulting residuals in nodal modular variability were used for subsequent analysis. Next, we selected the features using the training set by calculating the across-participant Pearson’s correlation between the modular variability of each node and the SRS score. Brain nodes showing significant correlation coefficients (i.e., *p* < .01) were retained. Thirdly, we built an SVR model to fit the relationship between the features (i.e., modular variability of selected nodes) and the SRS scores in the training set. Finally, we used the trained SVR model to predict the SRS score of the unseen participant (i.e., the test set).

After all LOOCV folds were completed, we assessed the performance of the prediction model by calculating the correlation between predicted and actual SRS scores. Then, we performed a permutation test (n = 10,000) by shuffling individual SRS scores to assess the statistical significance of the estimated prediction accuracy. Regional contribution to the prediction was defined as the frequency that each region was selected as a feature in the leave-one-out cross-validation. The SVR model was computed using the LIBSVM toolbox for MATLAB, with default settings (17) (https://www.csie.ntu.edu.tw/~cjlin/libsvm/).

#### Association between Alterations in Connectome Dynamics and Gene Expression Profiles

##### Estimation of Gene Expression in Brain Nodes

We utilized a brain-wide transcriptomic dataset of the adult human brain from the Allen Human Brain Atlas (AHBA) (18, 19) (http://human.brain-map.org), which includes post-mortem brain samples from six donors (i.e., five males and one female) aged 24-57 years. This dataset provides expression data for more than 20,000 genes in 3,702 spatially distributed tissue samples. Tissue samples from two donors cover the whole brain, while tissue samples from the other four donors cover only the left hemisphere. This dataset has been widely used to explore the genetic underpinnings of brain network properties in both typical development (20-23) and neuropsychiatric disorders (24, 25) due to the fact that canonical genetic signature of the adult human brain is highly consistent across individuals (19). Considering that rsfMRI data used here were obtained from only male subjects, we employed gene expression data of the five male donors to explore the genetic underpinnings of altered brain network dynamics in ASD.

AHBA gene expression data was preprocessed using the default procedures of a state-of-the-art pipeline (26) (https://github.com/BMHLab/AHBAprocessing). This included probe-to-gene re-annotation, filtering of probes that did not exceed background noise, selecting the representative probe for each gene, assigning each sample to its nearest node in the random-512 parcellation, and correcting for both inter-sample and inter-subject variability. Several details in relation to the preprocessing should be noted. First, we performed probe-to-gene re-annotation based on the latest data from the National Center for Biotechnology Information (NCBI) using Re-annotator (27). Secondly, we performed the spatial assignment of tissue samples to brain parcellation nodes in native space. Specifically, we first segmented T1 images of each donor brain using tissue probability maps in MNI space as the reference for segmentation. We then spatially transformed the random-512 parcellation from MNI space to the native space by applying the inverse deformation field obtained from the T1 segmentation procedure. Tissue samples for each donor were then matched to the spatially nearest node based on donor-specific parcellation. Third, only gene expression data from the left hemisphere was used to ensure the reliability of results, since gene expression data for the right hemisphere was available from only two male donors. In the left hemisphere, 222 out of 236 nodes were spatially matched with expression data, resulting in a 222 ×10,145 matrix, which denoted the expression of 10,145 genes across 222 nodes.

##### Enrichment Analysis

After performing the partial least squares regression (PLS) analysis, we ranked the 10,145 genes according to their weights to the PLS1 component. Genes ranked at the top of this ordered gene list exhibited the strongest positive associations with the ASD-related alterations in modular variability, which were overexpressed in regions showing higher modular switching in ASD. Genes ranked at the bottom of this ordered gene list exhibited the strongest negative associations with ASD-related alterations in modular variability, which were overexpressed in regions showing lower modular switching in ASD.

To gain further understanding of the biological implications of the genes identified as being associated with case-control differences in modular variability, we undertook two different enrichment analyses. First, we used the Gorilla tool (28) (http://cbl-gorilla.cs.technion.ac.il/) to perform Gene Ontology(GO) enrichment analyses to explore functional implications of the identified genes. Gorilla is a widely used online tool that identifies GO terms that are significantly enriched at the top of the ordered gene list, without pre-assigning thresholded target gene sets and background sets (28). To identify the GO terms that were enriched at the bottom of our ordered gene list, we reversed the ordered gene list and performed the GO enrichment analyses again. Second, we carried out a gene set enrichment analysis (GSEA) (29) on the whole gene list (i.e., the 10,145 ordered genes) to assess whether previously identified ASD-related gene sets were enriched at the top or bottom of our ordered gene list. The GSEA analysis does not require thresholding of the ordered gene list, which may reduce the effects of arbitrary thresholding and capture genes that may not show significant correlation with modular variability but nonetheless have important biomedical significance (29). Six classes of ASD-related gene sets and one gene set that was associated with non-mental health diseases were used in the GSEA. The six ASD-related gene sets are listed below:

i. Gene set 1: ASD-related gene set from a summary of multiple datasets (n = 594) (30). This gene set was curated by Krishnan et al, and was collected from both publicly available databases (e.g., SFARI Gene, http://gene.sfari.org) and automatically text-mined ASD-gene co-occurrences in published abstracts (e.g., Gene2Mesh, http://gene2mesh.ncibi.org/).
ii. Gene set 2: ASD risk genes identified from a large-scale exome sequencing study (n = 102) (31). Of these genes, 53 show higher frequency in individuals ascertained to have ASD, and the remainder show higher frequencies of disruptive de novo variants in individuals ascertained to have severe neurodevelopmental delay.
iii. Gene set 3: ASD-associated common genetic variants identified from a genome-wide association study (GWAS) meta-analysis study (n = 88) (32). We extracted this gene set from the summary statistical results of the GWAS meta-analysis study (32). Specifically, SNP-based *p* values were converted to gene-based *p* values using H-MAGMA (33). We selected genes with an FDR corrected *p* < .1 (33), which included 88 genes.
iv. Gene set 4: ASD associated rare, de novo variants identified from a study integrating copy number variants and sequencing data (n = 65) (34). A total of 65 ASD risk genes (FDR corrected *p* < .1) were identified by Sanders et al (34)
v. Gene set 5: genes upregulated in the ASD cortex, identified from a post-mortem genome-wide transcriptome study (n = 584) (35). We selected differential expression genes with FDR corrected *p* < .05. Upregulated genes were identified as those with log_2_(Fold-change) > 0 (36).
vi. Gene set 6: genes downregulated in the ASD cortex, identified from a post-mortem genome-wide transcriptome study (n = 558) (35). We selected differential expression genes with FDR corrected *p* < .05. Downregulated genes were identified as those with log_2_(Fold-change) < 0 (36).
vii. Gene set 7: a gene set associated with non-mental health diseases includes 1,189 genes (30), and was used to assess whether the PLS-derived genes were mental disease-specific.

GSEA was performed using the GSEA function in clusterProfiler package version 3.14.3 (https://bioconductor.org/packages/release/bioc/html/clusterProfiler.html) (37). The results were visualized using the gseaplot2 function in enrichplot package version 1.6.1 (https://bioconductor.org/packages/release/bioc/html/enrichplot.html) in R language.

#### Validation Analysis

We validated our main findings by considering five potential confounding factors. *(i) Head motion*. During image preprocessing for the validation analysis, an additional spike regression-based scrubbing strategy was employed during the nuisance regression procedure to further control the influence of head motion (38). “Bad” volumes were defined as those with framewise displacement above 0.5 mm and their adjacent volumes (one forward and two backward). *(ii) Window length*. In addition to the window length of 60 seconds used in the main analysis, we also reconstructed the dynamic functional networks with a longer window length of 100 seconds to assess the effect of the window length on our results. *(iii) Intelligence quotient (IQ)*. As part of the validation analysis, we included the intelligence quotient in the GAM as an additional covariate to reduce its influence on group differences. *(iv) Imaging sites*. We utilized a leave-one-site-out cross validation strategy to estimate whether group differences were biased by specific sites. *(v) Age range*. We reexamined the case-control differences by excluding participants aged over 21 years (39) to assess whether group differences were observable for the children and adolescents.

## Supplementary Results

### Age and Interaction Effects in the Mega-Analysis

At global level, the GAM-based analysis revealed a significant nonlinear age effect (i.e., a slight increase followed by a persistent decline) on modularity (*p* = .0009) (Figure S2A), but no group and age-by-group interaction effects (both *p*s > .05). In addition to the significant group effects on the mean and standard deviation of whole brain modular variability, we also observed a nonlinear age effect (i.e., a slight decline followed by a slight increase) on mean modular variability (*p* = .037) (Figure S2B), but not on standard deviation (*p* > .05). No interaction effect was observed for either metric.

At nodal level, in addition to the significant group effect on the modular variability at several default-mode and visual areas, we also identified significant age effects on modular variability, primarily in several frontal and temporal areas (*p* < .05, FDR corrected) (Figure S3A) such as the medial superior frontal cortex, supplementary motor area, superior temporal gyrus, and insular cortex. Only a few nodes showed significant age-by-group interaction effects, including the bilateral superior temporal gyrus and the right fusiform (*p* < .05, FDR corrected) (Figure S3B).

### Validation Results

When assessing the potential influence of head motion, window length, IQ, data collection sites, and age range, we found that the resultant maps of significant ASD-related alterations in modular variability remained highly similar to those in our main findings (Dice index range: .77 to .95) (Table S2 and Figure S5). This suggests that the spatial distribution of case-control differences identified in our main findings was highly robust and weakly affected by methodological variations. It is also worth noting that significant age-by-group interaction effects varied substantially between different validation strategies (Dice index range: 0 to 1), suggesting limitations in robustness. Thus, the findings regarding the interaction effects should be interpreted with caution.

### Supplementary Tables

**Table S1.** Number of overlapped genes between six ASD-related gene sets

|  | Gene set 1 | Gene set 2 | Gene set 3 | Gene set 4 | Gene set 5 | Gene set 6 |
| --- | --- | --- | --- | --- | --- | --- |
| Gene set 1 | 594 |  |  |  |  |  |
| Gene set 2 | 26 | 102 |  |  |  |  |
| Gene set 3 | 2 | 1 | 88 |  |  |  |
| Gene set 4 | 20 | 38 | 1 | 65 |  |  |
| Gene set 5 | 21 | 0 | 0 | 0 | 584 |  |
| Gene set 6 | 21 | 3 | 0 | 1 | 0 | 558 |

Diagonal elements denote the total number of genes in each gene set and the lower trigangular elements dennote the number of genes that overlap between each pair of gene sets. The gene sets were obtained from the following sources: i) gene set 1, ASD-related genes from a summary of multiple datasets (30); ii) gene set 2, ASD risk genes from a large-scale exome sequencing study (31); iii) gene set 3, ASD-associated common genetic variants from a GWAS meta-analysis study (32); iv) gene set 4, ASD associated rare, de novo variants from a study integrating copy number variants and sequencing data (34); v) gene set 5, genes upregulated in the ASD cortex from a post-mortem genome-wide transcriptome study (35); vi) gene set 6, genes downregulated in the ASD cortex from a post-mortem genome-wide transcriptome study. GWAS, genome-wide association study.

**Table S2.** Dice indices of thresholded statistical maps between the main results and different validations

| **Validation Strategy** | **Group effect** | **Age effect** | **Interaction effect** |
| --- | --- | --- | --- |
| IQ regression | 0.94 | 1 | 1 |
| Head motion | 0.77 | 0.77 | 0.33 |
| Window length | 0.80 | 0.75 | 0.67 |
| Age range | 0.87 | 0.58 | 0 |

Here, we assessed the robustness of the group, age and age-by-group interaction effects at the nodal level under four different validation conditions. “IQ regression” denotes the inclusion of the IQ term in the GAM to mitigate its influence. “Head motion” denotes the use of a spike regression-based scrubbing strategy in nuisance regression during image preprocessing to further control the influence of head motion, “Window length” denotes the reconstruction of the dynamic functional networks using a window length of 100 seconds, and “Age range” denotes excluding adult participants with ages over 21 years. Each value in the table represents the Dice index between the original thresholded modular variability map in the main findings and that obtained under a validation condition. GAM, generalized additive model; IQ, intelligence quotient.

## Supplementary Figures

**Figure S1.**
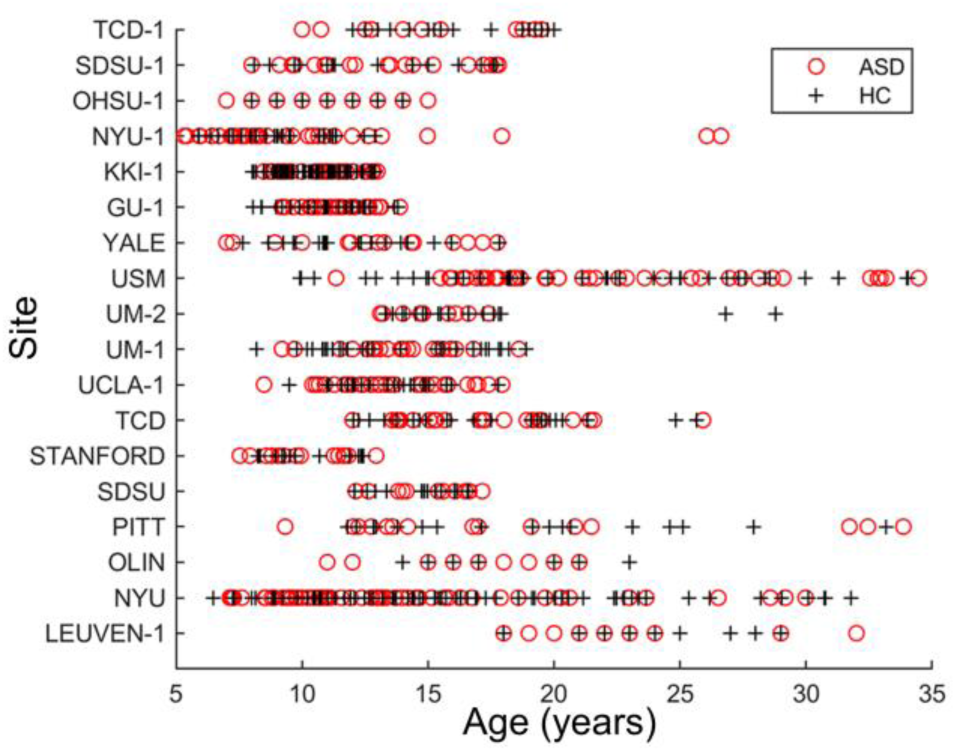
Distribution of participants across the 18 sites. For each site, both ASD and HC groups contained at least 10 participants. Red circles denote ASD individuals and black crosses denote HCs. ASD, autism spectrum disorder; HC, healthy control; TCD-1, Trinity Centre for Health Sciences: Sample 1; SDSU-1, San Diego State University: Sample 1; NYU-1, NYU Langone Medical Center: Sample 1; KKI-1, Kennedy Krieger Institute; Sample 1; GU-1, Georgetown University: Sample 1; Yale, Yale School of Medicine; USM, Utah School of Medicine; UM-1, University of Michigan: Sample 1; UM-2, University of Michigan: Sample 2; UCLA-1, University of California, Los Angeles; TCD, Trinity Center for Health Sciences; STANFORD, Stanford University; SDSU, San Diego State University; PITT, University of Pittsburgh; OLIN, Olin, Institute of Living at Hartford Hospital; NYU, NYU Langone Medical Center; LEUVEN-1, University of Leven: Sample 1.

**Figure S2.**
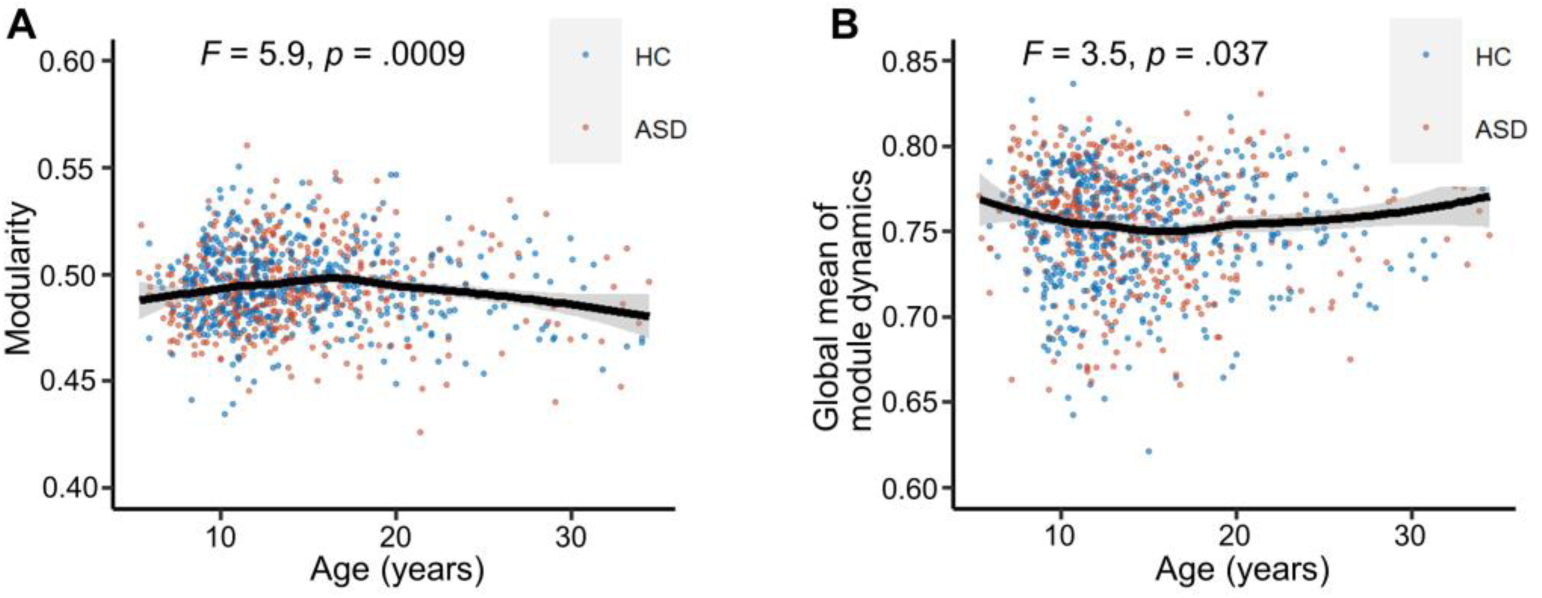
Age-related changes in global modular dynamics. (A) Significant nonlinear age effects on modularity. (B) Significant nonlinear age effects on the mean modular variability of the brain. We used a GAM to estimate age effects for the whole population, including both ASD individuals and HCs. ASD, autism spectrum disorder; HC, healthy control; MV, modular variability; GAM, generalized additive model.

**Figure S3.**
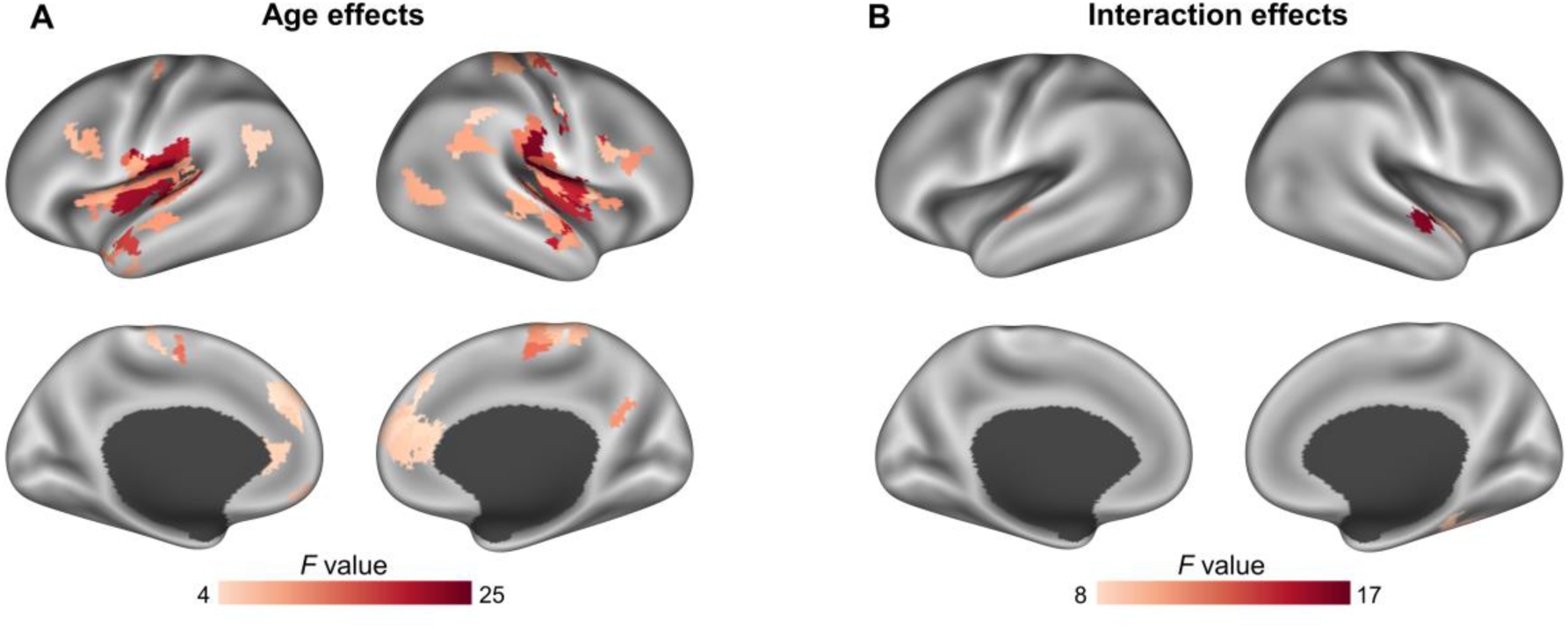
Age and age-by-group effects on regional module dynamics. (A) Age effects on nodal modular variability. (B) Age-by-group interaction effects on nodal modular variability. We used a GAM to estimate the age effects and the group-by-age interaction effects on nodal modular variability. Significance levels were corrected for multiple comparisons by pooling the *p* values of all three effects (i.e., group, age, and interaction) across all nodes. GAM, generalized additive model.

**Figure S4.**
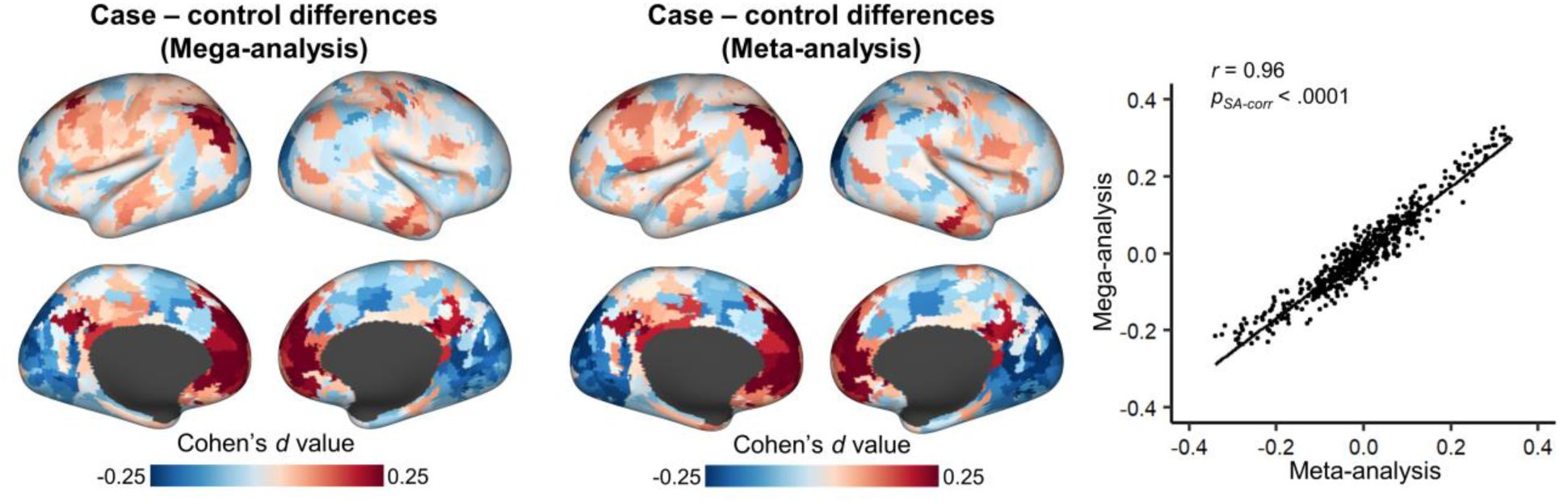
Spatial similarity between case-control difference maps obtained from the mega- and meta-analyses. The left and middle panels respectively represent the spatial pattern of case-control differences obtained from the mega- and meta-analysis. In the right-hand panel, each dot in the scatterplot represents a brain node. The significance level of the spatial association was corrected for spatial autocorrelation (40). SA-corr denotes correction for the spatial autocorrelation.

**Figure S5.**
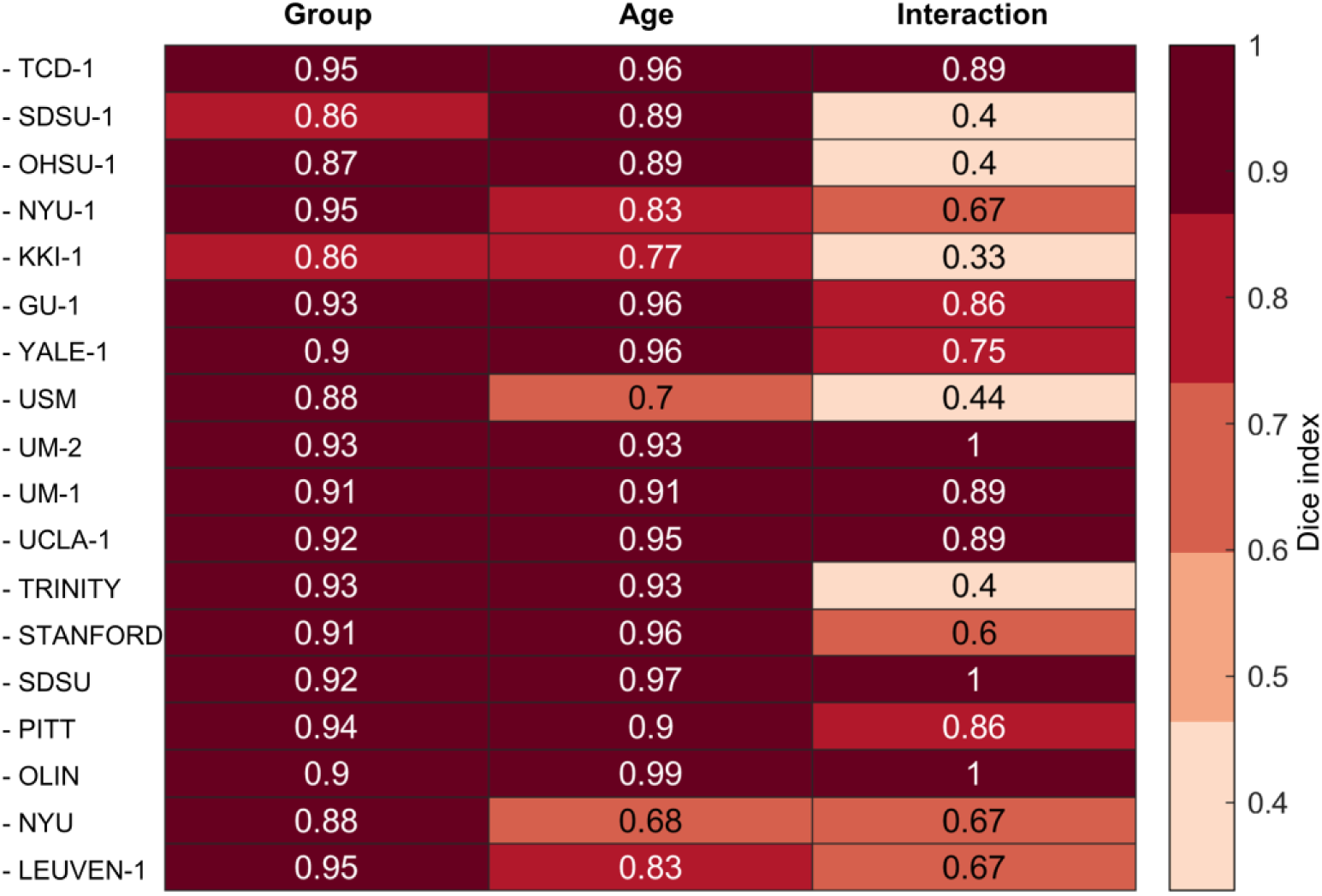
Dice indices between thresholded statistical maps in the main results and the leave-one-site-out cross-validations. We performed a leave-one-site-out cross-validation to assess whether our main findings were biased by site specific sites. Each label on the left represents the site being leaved out. Each column represents the effect of interest (i.e., group, age, and group-by-age interaction). Each number denotes the Dice index between the main results and the validation. TCD-1, Trinity Centre for Health Sciences: Sample 1; SDSU-1, San Diego State University: Sample 1; NYU-1, NYU Langone Medical Center: Sample 1; KKI-1, Kennedy Krieger Institute; Sample 1; GU-1, Georgetown University: Sample 1; Yale, Yale School of Medicine; USM, Utah School of Medicine; UM-1, University of Michigan: Sample 1; UM-2, University of Michigan: Sample 2; UCLA-1, University of California, Los Angeles; TCD, Trinity Center for Health Sciences; STANFORD, Stanford University; SDSU, San Diego State University; PITT, University of Pittsburgh; OLIN, Olin, Institute of Living at Hartford Hospital; NYU, NYU Langone Medical Center; LEUVEN-1, University of Leven: Sample 1.

## Notes

### Competing Interest Statement

The authors have declared no competing interest.

